# Assessing the digit organisation of focal hand dystonia using 7T functional MRI

**DOI:** 10.1101/2025.09.11.675579

**Authors:** Michael Asghar, Rosa Sanchez-Panchuelo, Paul Glover, Daisie Pakenham, George C. O’Neill, Denis Schluppeck, Miles Humberstone, Susan Francis

## Abstract

**Introduction:** Focal hand dystonia (FHD) is typically treated with Botulinum Toxin (BTX/BoNT-A/Botox/ Onabotulinum Toxin Type A) injected into digit and wrist muscles to provide temporary symptomatic relief from muscle spasm. We use 7T functional MRI (fMRI) to study 1) differences in somatosensory and motor cortical digit maps between FHD patients and age and sex-matched healthy volunteers (HV), 2) changes in cortical digit maps of FHD patients after BTX treatment.

**Methods:** Four FHD patients underwent behavioural measures and 7T fMRI at two timepoints: ∼30 days after BTX treatment at peak efficacy (Post-BTX), immediately prior to their next injection ∼100 days after treatment when therapeutic effects had declined (Pre-BTX). HVs were scanned at a single time point. Behavioural measures included temporal discrimination (TDT), amplitude thresholding (AMP) and spatial acuity grating orientation tasks. A somatosensory travelling wave (TW) digit-mapping fMRI task was performed on dominant/affected and non-dominant/un-affected hands, and a motor TW digit-mapping task on the dominant/affected hand. Fourier analysis derived digit maps which were compared to a probabilistic digit atlas, and population receptive field (pRF) mapping performed. Structural and resting state fMRI data were acquired.

**Results:** Spatial acuity was lowered in the dominant/affected hand in FHD compared to HVs, while TDT and AMP measures showed no significant differences. Somatosensory and motor TW cortical digit maps in FHD and HV showed clear digit ordering. Somatosensory and motor cortical digit maps were more disordered in the dominant/affected hand than the non-dominant hand for FHD Post and Pre-BTX compared to HVs (DICE coefficient/Figure of Merit). Compared to the atlas, HV maps were in-line for both hands, while in FHD central tendency (CT) was lower in the dominant/affected hand than non-dominant hand. pRF sizes were larger in the FHD Pre-BTX group compared to HV and Post-BTX for the dominant hand.

**Conclusion:** Behaviourally, FHD patients had lower spatial acuity than HVs. FHD digit maps had typical D1-D5 ordering, but DICE coefficients identified a disordered dominant hand in FHD patients compared to HVs at both timepoints. The FHD Pre-BTX group had larger pRF sizes than Post-BTX and HVs, suggesting BTX treatment coincides with more localised touch perception.

## 1. INTRODUCTION

Focal Hand Dystonia is a sensorimotor disorder presenting with involuntary movements, tremors and weakness in the hand (1). Early brain imaging studies of focal hand dystonia (FHD) have suggested disordered somatosensory function demonstrated by reduced distances between the cortical representation of digits and increased overlapping activity compared to healthy controls (2,3). However, because the cortical digit maps of somatosensation and motor function representing the hand are at the millimetre scale, these findings of disorder are now being questioned as higher spatial resolution fMRI become available, motivating this work.

Functional MRI (fMRI) studies probing the cortical digit representation use block (4,5), phase-encoding (6) and event-related (7) designs to deliver sensory stimulation to the fingers, mostly in healthy individuals. In the past decade, there has been a push to investigate digit maps at higher magnetic field strength, due to the increased BOLD contrast and spatial resolution (8,9). Somatotopic digit mapping in human primary somatosensory cortex using a travelling-wave (TW) phase-encoding fMRI paradigm has been shown to be highly reproducible in healthy participants at 7T (10) but show inter-participant variability. To address this, we developed a probabilistic somatosensory atlas (11) to define digit dominance and inter-participant variability in healthy participants. This atlas provides a standardised template for comparison with patient groups, such as those people with Focal Hand Dystonia (FHD).

Prior studies in dystonia have principally used motor tasks. A study in dystonic musicians (12) showed greater activity in ipsilateral premotor area and reduced activity in the cerebellum during a right-handed tapping task, suggesting irregular neural activity and compensatory mechanisms compared to healthy controls. In contrast, a study of a cued movement task in FHD patients (13) showed a decrease in ipsilateral activity and an increase in contralateral activity associated with the affected hand. A further study (14) showed that motor performance to a button press task was similar between controls and FHD patients, while the motor preparation section of the task showed reduced activation in premotor and postcentral gyrus for FHD patients, implying impaired early stages of motor control. A 3T fMRI study using passive extension and active finger presses of individual fingers in musician’s dystonia (n=9) (15) discerned individual digits and clear somatotopy but showed no difference in the spatial geometry (expansion or spatial shift of digit representation) for musicians with and without dystonia. This suggestion of the intact cortical digit maps in musician’s dystonia contests the idea that task-specific dystonia alters cortical maps and lends weight to an alternative hypothesis that task-specific dystonia is due to a higher-order disruption of skill encoding. A recent high spatial resolution (0.75×0.75×0.89mm) laminar 7T MRI Vascular Space Occupancy (VASO) study showed in a small group of individuals with FHD (n=4) a breakdown of digit somatotopy and increased activation in superficial cortico-cortical input layers during finger tapping compared to unaffected hemispheres, suggested to be due to reduced inhibition (16).

Together, these studies of FHD suggest widespread differences in sensorimotor areas and subcortical structures. However, these prior studies have generally been performed at a field strength of 3T or less and have used spatial analyses (e.g. centre of gravity of each digit based on vertices and searchlight analysis) in volumetric space. There is therefore a need for high spatial resolution high field fMRI studies using surface-based methods to provide more accurate and more sensitive results (17) and better digit delineation across gyral and sulcal boundaries. More recently, fMRI studies have measured how receptive fields of neurons in the somatosensory cortex are distributed, for example by using a 2D grid of stimulators to map within- and between-digit characteristics. Computational models have been used to investigate more fine-grained characteristics of the hand, such as population receptive field (pRF) size, in healthy participants (18–21).

Resting state fMRI (rs-fMRI) has also shown alterations in functional connectivity (FC) in dystonia. In Writer’s Cramp (WC) (22) reduced functional connectivity compared with controls was shown when placing seeds in left lateral premotor cortex, left thalamus, left/right pallidum to study the projection to the symptomatic left primary sensorimotor cortex. There were also reductions in FC from primary motor cortex premotor, frontal and somatosensory areas. Reductions in FC were also shown in cervical dystonia, in regions related to the M1-SMA motor network (23). Another study showed abnormal recruitment of parietal and premotor cortices, as well as distinct patterns of sensorimotor integrations between musician’s and non-musician’s dystonia (24). Unique patterns for dystonic tremor, based on a reduction in FC were found in higher-level cortical, basal ganglia, and cerebellar regions (25). Further, a study of cervical dystonia and blepharospasm (26) suggested that focal dystonia is a disorder of specific networks and not just a result of basal ganglia alterations.

Behaviourally, there is evidence of raised temporal thresholds in patients as determined with temporal discrimination tasks and spatial discrimination thresholds using a grating orientation task (GOT) (27,28).

A symptomatic treatment of FHD involves injections of Botulinum Toxin (BTX), into the muscles controlling digit and /or wrist movement. This can induce relief from involuntary muscle spasm associated with FHD. A prior fMRI study in patients with cervical dystonia affecting the neck showed an increase in activation in sensorimotor cortex, 4 weeks after BoNT injection (29).

Here we used high spatial resolution 7T fMRI and surface-based analyses to study cortical somatotopy digit maps and the pRF digit size and perform rs-fMRI in a group of FHD patients and healthy controls (HV). Patients were scanned at two timepoints in the BTX treatment cycle of, ∼30 days after localised BTX injection at peak efficacy (Post-BTX), and immediately prior to their next injection ∼100 days after treatment when therapeutic effects have declined (Pre-BTX). At each timepoint, behavioural tasks of temporal discrimination, amplitude thresholding and spatial acuity were collected.

We test the following hypotheses:

1. Behavioural thresholds (temporal and spatial) are raised in FHD patients compared to healthy controls.
2. In FHD, digit maps derived from TW mapping (11) are blurred/have poorer delineation compared to healthy controls,
3. pRF sizes are increased in FHD patients compared to healthy controls.
4. Functional connectivity (FC) to the somatosensory cortex is reduced in FHD compared to healthy controls.

For these measures we predict FHD responses at 4 weeks following BTX treatment are more like those of healthy controls compared those at 3 months following BTX treatment.

## 2. MATERIALS AND METHODS

The study was conducted with the approval of the University of Nottingham Faculty of Medicine and Health Sciences Ethics Committee for healthy volunteers and of the Health Research Authority Research Ethics Committee (REC 17/EM/0368) for FHD patients. FHD patients were recruited from the Botulinum Toxin Clinic at Queens Medical Centre. Eligibility criteria included patients 18 and 75 years who were clinically assessed to have FHD and presented with unilateral symptoms on their dominant hand. Exclusion criteria included participants with a history of neurological illness other than FHD and MRI contraindications. All participants gave written informed consent and completed a MR safety screening form and a handedness form at each visit.

Six patients (2 visits) with Focal Hand Dystonia (FHD) (age mean ± stdev: 54 ± 11 yrs, 4 male) were recruited to be scanned at 7T at both 4 weeks (20-27days) (Post-BTX) and 3 months (105-119 days) after (Pre-BTX) Onabotulinumtoxin type-A (Botox, BTX) treatment (30), along with age-matched healthy controls (HVs) (age mean ± stdev: 42 ± 7 yrs. At each visit, participants completed 1hour of behavioural data collection followed by 7T functional and structural MRI scanning lasting 1hour. In all FHD participants, the affected hand was also the dominant hand.

### 2.1 Behavioural data acquisition

Behavioural data collection included a temporal discrimination task (“TDT”) and an amplitude threshold task (“AMP”) coded in MATLAB (The Mathworks, Natick, Massachusetts), and spatial acuity grating orientation task (“GOT”). These tasks were performed using identical piezoelectric stimulators to those used in the fMRI study, while the GOT used Perspex domes.

#### TDT

The temporal discrimination task was performed on the tips of digits D2 and D3 of both the left and right hands using piezo stimulators, as well as a Brain Gauge device (31). On each trial, two brief stimuli were presented sequentially to D2 and D3, and participants had to indicate the digit they felt the stimulus appear first verbally; the experimenter noted down their responses. To estimate each participant’s temporal discrimination threshold, we used an adaptive 1-up, 2-down staircase procedure implemented via the *upDownStaircase()* function from MGL (32). This staircase method converges on the stimulus level at which participants respond correctly 70.7% of the time (33). The staircase ended after 8 reversals (changes in staircase direction).

#### AMP

To determine, participants completed a two-interval forced choice task to estimate on the left and right index finger (D2) at 30 Hz (match fMRI paradigm) and 200 Hz. On each trial, participants were presented with two brief tactile stimuli in succession: One stimulus was at baseline, near-threshold amplitude, the other was a test stimulus at higher amplitude, adjusted trial-by-trial via a 1-up, 2-down staircase procedure. The order of the baseline and test stimulus was randomized across trials. Participants were asked to indicate which of the two stimuli felt stronger. The task consisted of 60 trials, and the threshold estimate was taken as the mean amplitude across the final reversals. This procedure adaptively estimated the smallest amplitude difference the participant could reliably detect above baseline.

#### GOT

This grating orientation task was used to assess spatial acuity (34) of the tip of D2, due to time constraints only one hand was assessed: for FHD patients the affected hand was studied, and for controls the dominant hand. Participants were presented with machine-cut Perspex domes, each with a raised grating surface of a specific spatial frequency (defined by the groove-to-groove gap width in mm). On each trial, the experimenter applied a dome to the fingertip in one of two orientations: 0°: aligned with the long axis of the finger (“forwards”), 90°: perpendicular to the long axis (“sideways”). The participant, unable to see the dome, was asked to report whether the grating was oriented along (0°) or across (90°) the digit, by saying “forwards” or “sideways”. Dome orientation was randomised on each trial using MATLAB code. Each grating width was tested in 20 trials (10 trials per orientation), and orientations were pseudorandomised within each spatial frequency. Accuracy was recorded manually. Different stimulus sets were used for patient and controls to ensure a suitable range for estimating threshold performance in both groups (Controls: 0.75, 1.0, 1.2, 1.5, 1.75, 2.0, 2.5, 3.0 mm; 2) Patients: 0.75, 1.0, 1.5, 1.75, 2.0, 2.5, 3.0, 4.0 mm), since patients were unable to reliably discriminate orientation even at larger widths.

In addition, 10 healthy participants (age mean ± stdev: 36 ± 11 yrs) were recruited to assess the repeatability of the behavioural tasks. The participants underwent the same procedure as those in the main study (TDT using piezo, AMP and GOT). The tasks were each performed twice, separated by a two-week interval. This allowed both the within- and between-participant coefficient of variation (CV) of measures to be computed, and the comparison of the CV to (Mikkelsen et al. 2020) (35) who assessed reproducibility of vibrotactile detection and discrimination.

### 2.2 MRI data acquisition

Data were collected on a 7 T Philips Achieva scanner using a 32-channel receive coil (Nova Medical). Travelling wave (TW) fMRI data was collected to assess somatotopy and motortopy. Somatosensory/motortopy fMRI scans were collected using an axial 48 slice GE-EPI BOLD acquisition with TE/TR = 25/2000 ms, SENSE 1.5, 1.5 mm isotropic resolution collected using multiband (MB) factor 3.

Somatosensory stimuli were delivered to the fingertips (D1/D2/D3/D4/D5) using MR-compatible piezo-electric stimulators driven at 30 Hz vibrotactile frequency (Dancer Design, UK; http://www.dancerdesign.co.uk/). Stimulation was on for 4s per digit, with 20s per cycle, for 8 cycles. This was performed in the forward and reverse direction on each hand. (10). Somatotopic maps were generated for both the left and right hands in each scan session (Post-BTX and Pre-BTX for FHD patients).

Motortopy was performed in the dominant hand. During the motortopy task, participants were visually cued by 5 blinking (1Hz) circles to perform in-air tapping with the same timings as the somatosensory TW task. During the motortopy task, an accelerometer glove, designed in-house to monitor 3-axes of displacement, was attached to the participant’s dominant hand and used to track the precise timings of the in-air tapping for each of the five digits.

Resting state fMRI data was collected comprising 72 axial slices centred on the fMRI acquisition stack with a TE/TR=25/1500 ms, SENSE 1.5, 1.5mm isotropic resolution, using MB factor 4.

Structural data comprised a whole head T_1_-weighted MPRAGE (1 mm isotropic resolution) and PSIR (0.7 mm isotropic resolution), and high-resolution (0.5×0.5×1.5 mm^3^) T_2_*-weighted FLASH image, these were all collected with the same slice prescription as the fMRI data.

### 2.3 MRI analysis

Following quality control (tSNR, movement <1mm), only four participants were used going forward. fMRI data were distortion-corrected (FSL TOPUP), motion-corrected and aligned to the FLASH image. Data were high-pass filtered with a 0.01 Hz cutoff (100 s) to remove low-frequency trends. fMRI data were then analysed using custom software within mrTools (36) implemented in MATLAB. We performed several analyses to study the spatial representation of the digits and compare FHD patients to healthy controls, as described in Sections 2.3.1-2.3.5.

For these analyses, tissue segmentation and cortical reconstruction of the T_1_-weighted volumes was performed using Freesurfer (http://surfer.nmr.mgh.harvard.edu/) (37). Reconstructed cortical surfaces were flattened in a patch around the hand representation in post-central gyrus (radius, 55mm to 75 mm, depending on anatomy), using the mrFlatMesh algorithm (Vista software, https://github.com/vistalab/vistasoft/tree/master/mrAnatomy/mrFlatMesh). Freesurfer was used to estimate, for each participant, the location of primary and secondary sensorimotor Brodmann areas (BAs 1, 2, 3a, 3b) based on the histological analysis (38). The maximum-probability map for these was imported into participant-space to define ROIs for each Brodmann area. Data were registered to the *fsaverage* template (Freesurfer).

We also confirmed our digit representations from phase analysis of the TW data against a General Linear Model (GLM) analysis to confirm correspondence between analysis methods (see **Supplementary Methods).**

#### 2.3.1 Somatotopy and motortopy phase analysis of TW data

Somatotopy and motortopy data were analyzed using standard, Fourier based methods (in mrTools), producing coherence and phase maps. To account for shifts in the fMRI response due to the haemodynamic lag, the forward and reverse scans were averaged, as described in (7). Coherence, phase, and amplitude of the best-fitting sinusoid were then computed as described in (39). The phase map was displayed on the flattened cortical patch with a threshold of c>0.3 (coherence); this corresponds to an uncorrected p value of p<0.0004. First the digit correspondence between the somatosensory task data and the probabilistic Atlas were compared using Dice coefficients in fsaverage space for healthy volunteers and FHD patients for each visit. Then digit correspondence of the somatosensory and motor data was assessed using Dice coefficients between the HVs and FHD patients for each visit (Post-BTX and Pre-BTX), and between FHD Post-BTX and Pre-BTX. For each group, we also assessed Dice coefficients related to the extent of correspondence of motor and somatosensory maps in the Dominant hand. For the comparison between HV and FHD patients the Figure of Merit (FoM) was calculated from the Dice scores to summarise the response of each digit; this metric penalises degeneracy and poor representation (40).

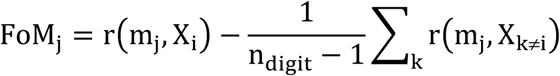

which defines the Dice (r) of the participant’s digit ROI (m) to its paired ROI (X) subtracting the mean of the Dice from the other digit ROIs. A FoM score of 1 represents perfect overlap with the target digit, whilst a lower score suggests the digit overlaps with the non-target digit or the representation is degenerate (i.e. overlaps with many digits).

#### 2.3.2 Comparison of data to the somatosensory probabilistic atlas

ROIs of each digit from the TW phase analysis were normalised to standard-space (*fsaverage*) and compared to the somatosensory probabilistic atlas (11). Surface projection and normalisation followed the methods described in O’Neill *et al*., i.e., using Freesurfer’s *mri_vol2surf* to project coherence maps, phase maps, and digit ROIs into the participant’s surface space, followed by a surface-to-surface transformation to MNI-305 space with *mri_surf2surf*. Since surface-based registration has previously been shown to outperform volume comparisons (11), only surface-based transformations were performed in this work. Full probabilistic maps (FPMs) were defined for each group (HC, and FHD for both Post-BTX and Pre-BTX), in which probabilities at each vertex were defined as the number of participants for which a particular digit was assigned to this voxel/vertex, divided by the total number of participants.

For the somatosensory data, central tendency (CT) was measured between the FHD patients’ digits and the atlas digits. CT defines how centrally located a digit is relative to a probabilistic atlas digit (1=perfect overlap, >1=resides centrally, <1=resides peripherally) (41). CT quantifies how much a single participant’s digit ROI overlaps with high probability areas of the atlas. For example, a small focal ROI that is within the atlas ROI would give a high central tendency. A larger ROI that encompasses the atlas ROI would also give a large central tendency if the two centres were aligned.

#### 2.3.3 Population receptive field (pRF) mapping

This analysis followed the methods described in (21) for somatosensory and motortopy tasks, which was adapted from a pRF method developed for visual cortex (42). Briefly, a 1-dimensional Gaussian model was combined with the spatio-temporal description of the stimulus, producing a predicted “cortical” response. To account for haemodynamic effects, this was then convolved with a HRF and then fit to the measured timeseries using non-linear least squares optimisation. This method produces cortical maps of “between digit” (equivalent to the TW Fourier analysis, albeit parameterized), pRF “size” (defined from the standard deviation of the 1D Gaussian best fit for each voxel) and adjusted r^2^ (the goodness of fit of the pRF model to the fMRI timeseries, pRF outputs were threshold by adjusted r^2^ > 0 (since adjusted r^2^ can be negative). ROIs were calculated by intersecting the binned digit phase value (D1-D5) with the FreeSurfer Brodmann areas (3a,3b,1,2), creating 20 ROIs for each participant. pRF size was extracted from each ROI and normalised (converted to z-scores) to account for inter-participant variability in absolute pRF size.

#### 2.3.4 Resting state connectivity

Data were analysed using the CONN toolbox in MATLAB (RRID:SCR_009550) (43) (Whitfield-Gabrieli and Nieto-Castanon 2012). First, the default preprocessing pipeline was used which included realignment, outlier detection, co-registration, normalisation to the MNI template, and 5 mm spatial smoothing. Denoising was performed using aCompCor (44), scrubbing, and temporal band pass filtering [0.008 0.09Hz]. Seed-based functional connectivity (FC) measures to cortical brain regions were then assessed by placing seed ROIs in both the left (L) and right (R) post-central gyrus taken from the Harvard-Oxford atlas (“atlas.PostCG r” and “atlas.PostCG l”). FC was represented by Fisher-transformed bivariate correlation coefficients from a weighted GLM, defined separately for each pair of seed and target areas, modelling the association between their BOLD signal timeseries. Cluster-level inferences were based on parametric statistics from Gaussian Random Field Theory; results were threshold using a cluster-forming p<0.001 voxel-level threshold, and familywise corrected p-FDR < 0.05 cluster-size threshold. Group independent component analysis (groupICA) was performed. FC was compared for 1) HC vs FHD (for both Post-BTX and Pre-BTX), and 2) FHD between visits (Post-BTX vs Pre-BTX).

##### Data and code availability statement

Code to generate results and figures, and the accompanying derived data is freely available on the Mendeley data repository: https://data.mendeley.com/datasets/tvt7mg5bf3/1.

## 3. RESULTS

### 3.1 Behavioural measures

Spatial acuity using the “GOT” task was significantly reduced in FHD for both Post-BTX and Pre-BTX groups compared with healthy controls (Mean HV (std): 1.87 (0.19), Mean Post-BTX (std): 2.96 (0.93), Mean Pre-BTX (std): 3 (0.88); one-way ANOVA p<0.05), **Figure 1A**.

**Figure 1:**
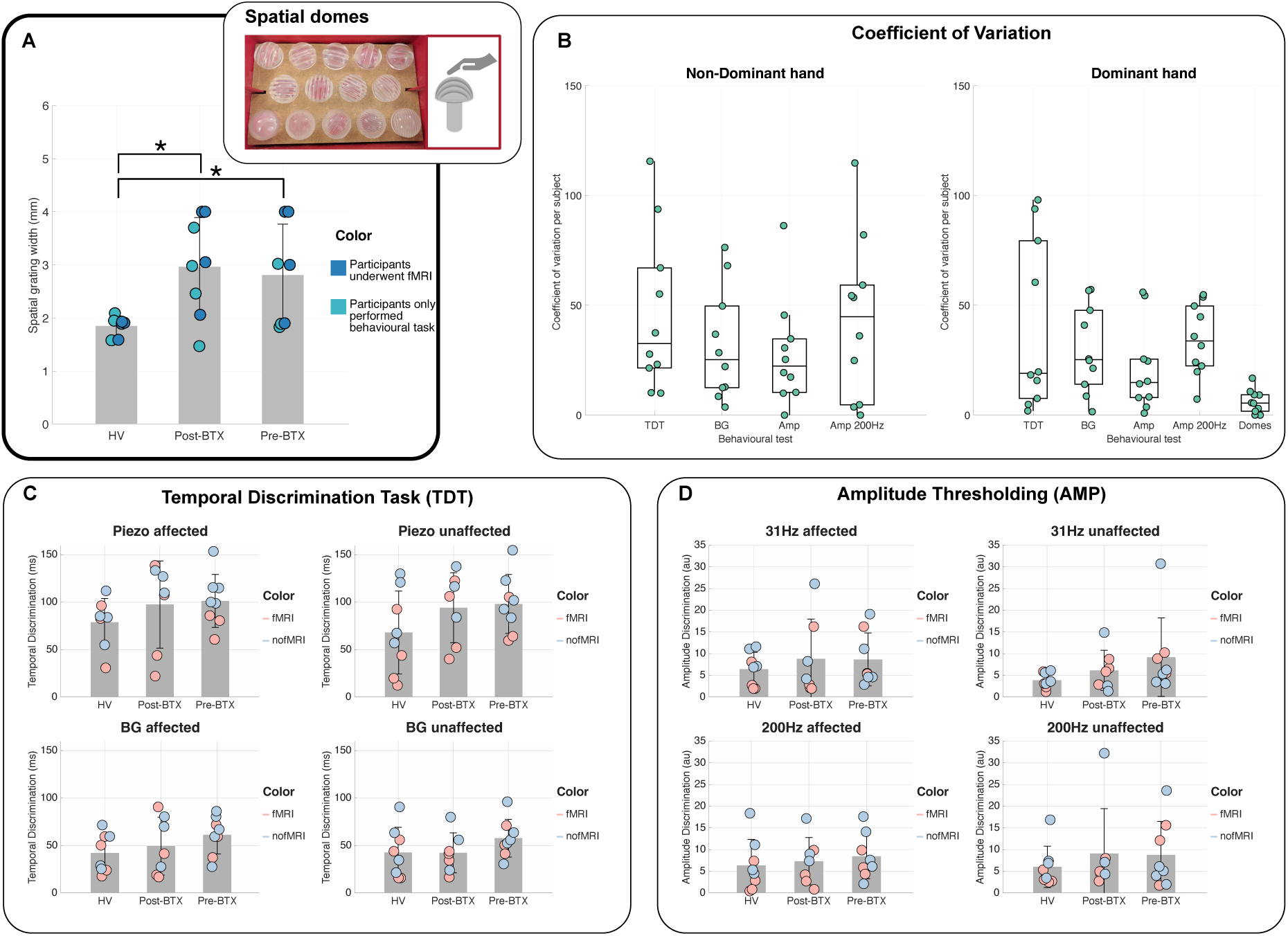
Behavioural results in healthy volunteer and focal hand dystonia Post-BTX and Pre-BTX visits for A) Spatial acuity grating orientation task (“GOT”) applied to the tip of the dominant (affected) hand. Participants who only performed the behavioural tasks and did not proceed to do the fMRI session are colour coded. B) Within-participant coefficient of variation (CV) computed from two visits in 10 healthy controls. CVs are comparable to Mikkelsen et al. 2020 (35); C) Temporal discrimination task (“TDT”) using piezoelectric stimulators and Brain Gauge (BG), and D) Amplitude discrimination (“AMP”) at 31 and 200 Hz.

In the healthy control repeatability group, within-participant CV for behavioural measures were: “GOT”: 6.2 ± 1.7 %; “TDT”: 40 ± 12 % (R) 46 ± 11 % (L) (Mikkelson: 46%); “AMP”: 21± 6 % (R) 28±8 % (L) (Mikkelson: 31%), as shown in **Figure 1B** and **Supplementary Table 1** which provides between-participant CVs. These CVs are in-line with those reported by Mikkelson (35).

There were no significant differences in the “TDT” tasks or “AMP” task between the FHD group and healthy controls, or for FHD patients between Post-BTX and Pre-BTX visits (**Figure 1C&D)**.

### 3.2 Comparing Travelling Wave digit maps between HV and FHD patients

Qualitatively, FHD travelling wave somatotopy and motor phase analysis digit maps showed similar digit-specific activity patterns to HVs, with FHD patterns similar between Post-BTX and Pre-BTX visits, **Figure 2**.

**Figure 2:**
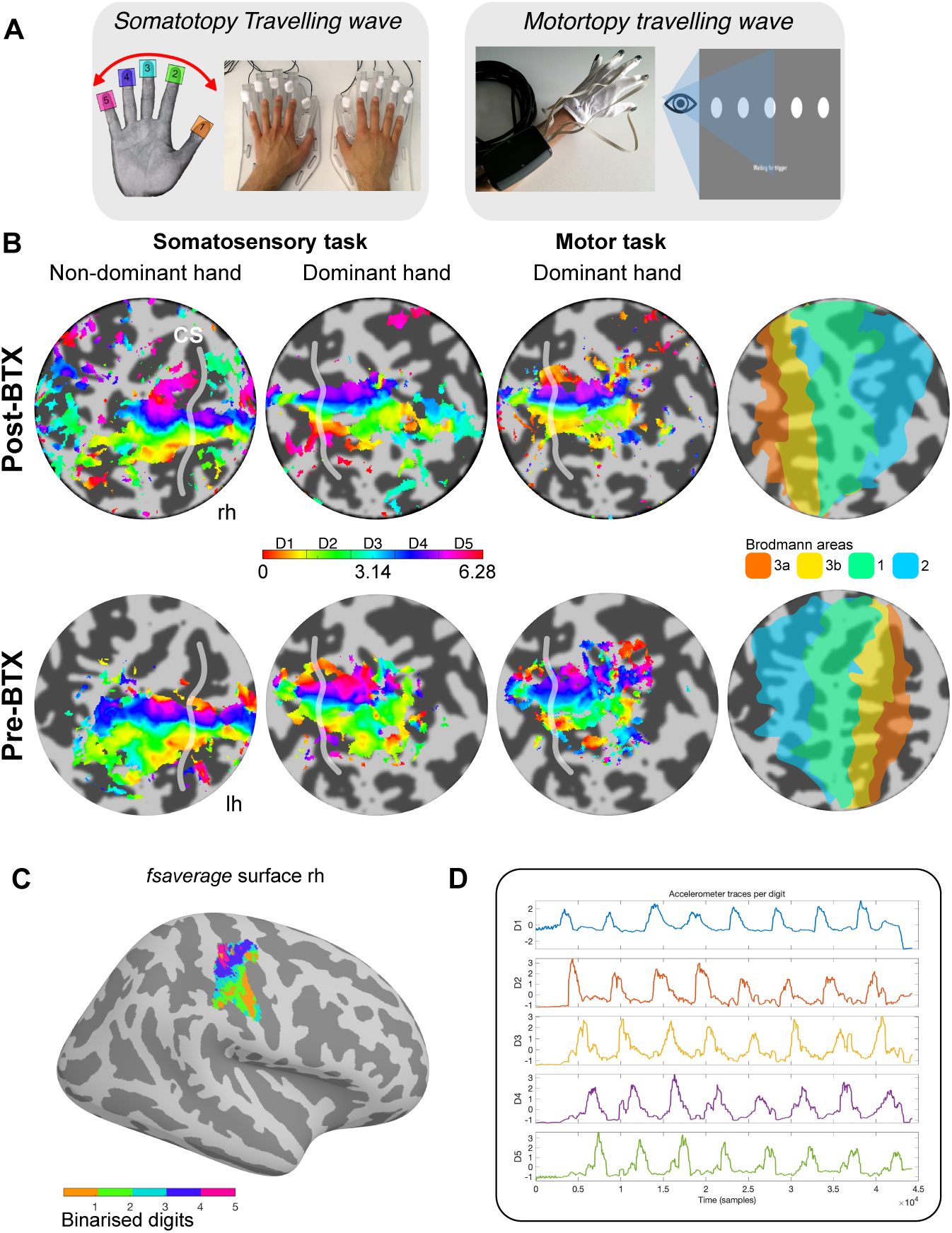
A) Somatosensory and motor travelling wave (TW) tasks performed during fMRI. An accelerometer glove was worn during the motor task to quality control participant movements. B) Example TW phase analysis digit maps to the Somatosensory and Motor task for an FHD participant at Post-BTX and Pre-BTX visits. Brodmann areas projected from the FreeSurfer segmentation also shown. C) Non-dominant somatosensory task TW map from Post-BTX visit projected to ‘fsaverage’ space. D) Example accelerometer trace from the forward run, post-BTX, dominant hand of the participant shown in B&C) for the motor task.

Quantitatively, Dice coefficient matrices of somatosensory phase analysis digit correspondence assessed in *fsaverage* space between groups are shown in **Figure 3**. When comparing matrices between HV and FHD groups for the somatosensory task (**Figure 3Ai**) and motor task (**Figure 3Aii**), lower Dice coefficients were seen in the dominant affected hand compared to the non-dominant hand for HV versus Post-BTX and HV versus Pre-BTX (P<0.02). This is shown in **Figure 3Aiv**) by the lower FoM of the Dice scores in the Dominant affected hand. However, no significant difference in the Dice matrices was detected between Post-BTX and Pre-BTX conditions. **Figure 3Aiii** is a graphic that explains how the DICE matrix affects the FoM score.

**Figure 3:**
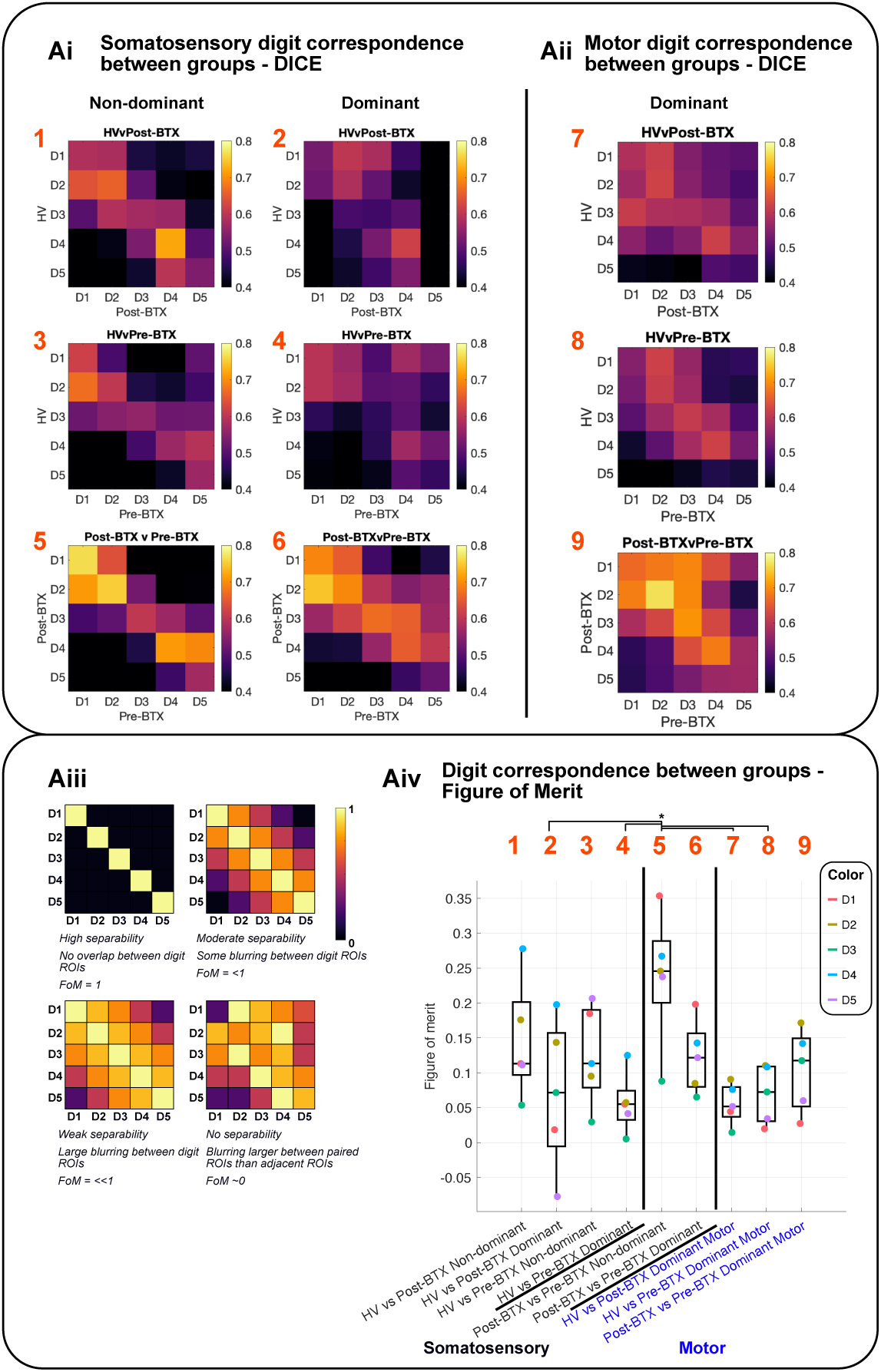
Ai) Dice matrices of digit patterns between study groups (HV versus FHD Post-BTX visit, HV versus FHD Pre-BTX visit and FHD BTX versus NoBTX) for (i) somatosensory task for the Non-dominant (left, non-affected) and Dominant (right, affected) hand and (ii) motor task for the Dominant hand. Aiii) Graphic to outline how key patterns and how the Figure of Merit (FoM) relates to the DICE matrices. Aiv) Figure of Merit (FoM) computed for each DICE matrix in (Ai) and (Aii). Note the FoM tends to be lower for the dominant hand. Significant difference (p<0.05, corrected for multiple comparisons) in FoM shown by brackets above plot.

In response to the somatosensory task the dominant hand had larger digit ROIs than non-dominant digit ROIs for FHD and HV groups (P<0.03), with D5 being significantly smaller than all other digit (D1-D4) ROIs (P=0.004) (see **Supplementary Figure 1**).

Somatosensory and motortopy task dominant hand digit maps showed strong spatial correspondence, with clear diagonal high Dice coefficients in each HV, FHD Post-BTX, FHD Pre-BTX group, **Figure 4**. For all participants, there was good agreement in digit correspondence of the TW GLM “winner-take-all” maps with the standard phase analysis maps shown by the high diagonal Dice matrix coefficients (**Supplementary Figure 2**).

**Figure 4:**
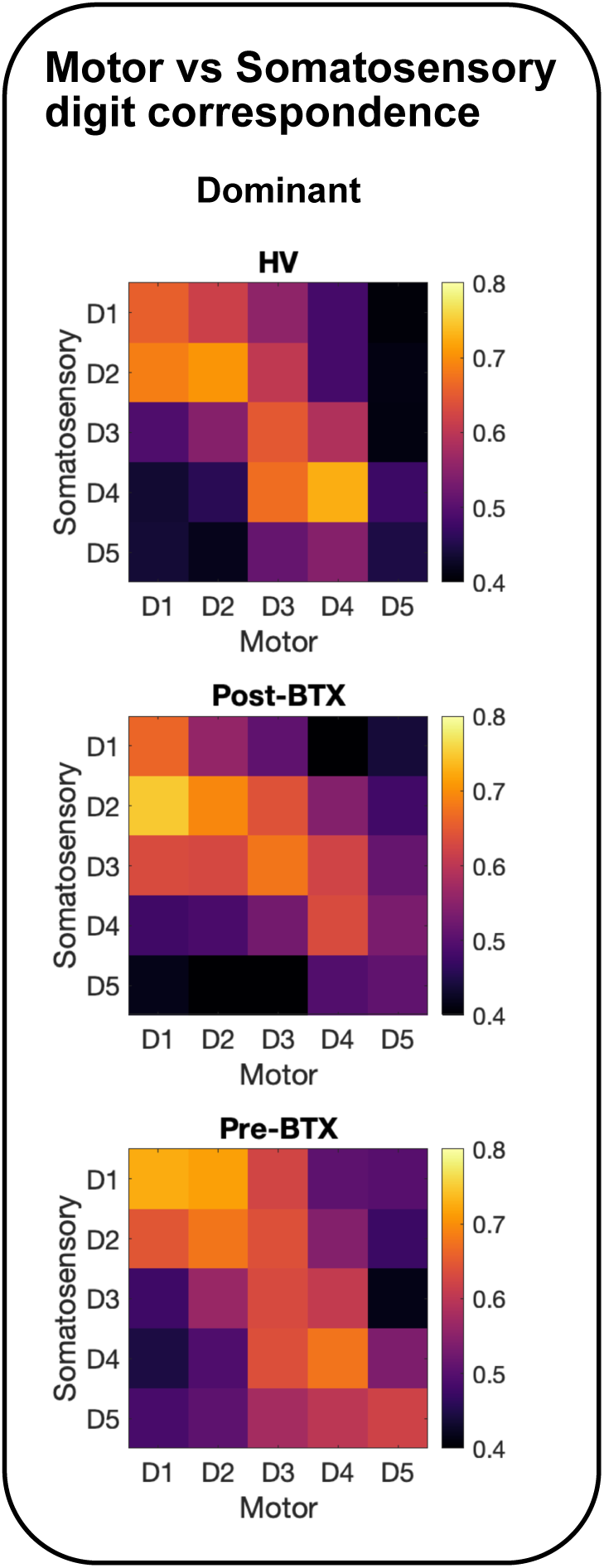
DICE matrices of digit patterns between motor and somatosensory maps for the Dominant (right, affected) hand for each study group. Strong correspondence is seen for the HVs and FHD patients at both visits.

### 3.3 Comparing digit maps to the probabilistic atlas

The Dice of the TW phase analysis digit maps in response to the somatosensory task with the Atlas digit ROIs showed a trend for a stronger diagonal in the HV compared to FHD group, particularly for the Dominant (affected) hand, **Figure 5A** (mean Dice diagonal: HV L/R: 0.634 ±0.009/0.62±0.02; Post-BTX L/R: 0.61 ±0.02/0.60±0.01; Pre-BTX L/R: 0.58 ±0.02/0.58±0.01). This is also reflected in the central tendency (CT) matrices (**Figure 5B**) which also considers how centrally located a digit is. A within group paired t-test between Dominant and Non-dominant hand CT scores, showed that for both FHD visits (Post-BTX and Pre-BTX) the Dominant hand had significantly smaller CT values than the non-Dominant hand (p<0.05, Bonferroni corrected).

**Figure 5:**
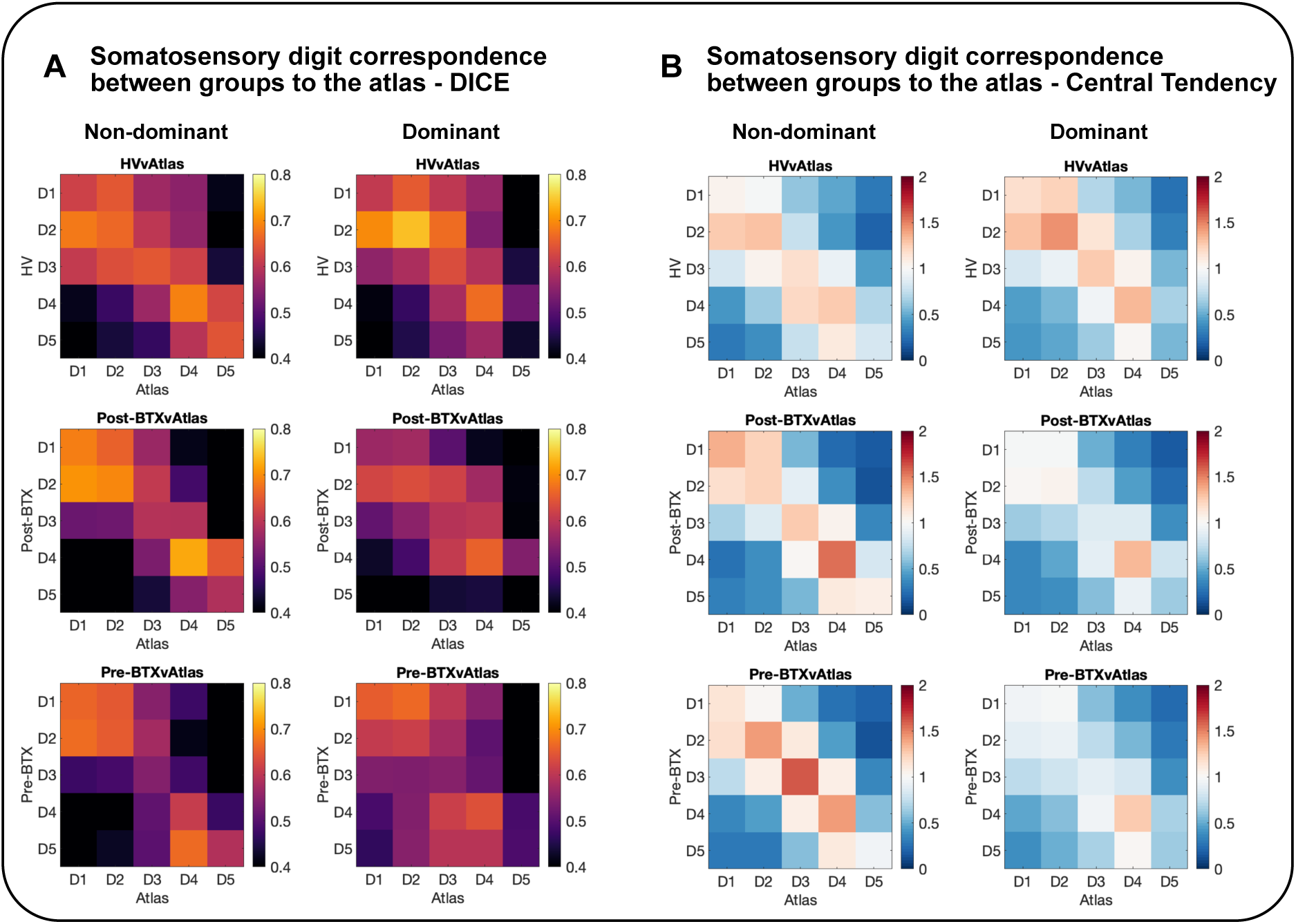
A) Dice matrices of digit patterns for the somatosensory task between HV, FHD groups compared with the probabilistic atlas. B) Matrices of central tendency (CT) scores for HV and FHD Post-BTX and Pre-BTX visits. The CT score represents how centrally located the digits are to the probabilistic atlas (1=perfect overlap, >1=resides centrally, <1=resides peripherally). Note no significant difference in CT matrix between Non-dominant and dominant hand for HV vs Atlas, while there was a significant difference (P<0.001) in CT matrix between Non-dominant and dominant hand for FHD vs Atlas for both Post-BTX and Pre-BTX visits.

### 3.4 Population Receptive Field (pRF) mapping

Mean (absolute) pRF size (across hemispheres and digits) across groups was HV (1.7±1.8), HV Atlas (1.4±1.2), FHD Post-BTX (2.0±1.9), FHD Pre-BTX (1.7±1.9), these results are in-line with our previous study in HVs (1.9±1.6 2D Gaussian model, mean±stdev) (21).

Comparing group mean pRF sizes for the somatosensory task, the FHD Pre-BTX group had larger pRF sizes in the Dominant (affected) hand than the FHD Post-BTX, HV, and Atlas. There were no differences in pRF size between groups in the Non-dominant hand for the somatosensory task, or in the motor pRF data for the Dominant hand (**Figure 6**). There were no differences in pRF sizes from the motortopy data in the Dominant hand (the only hand on which the motor task was performed). There was no overall significant difference in pRF size between hands across groups. Motor pRF size was larger than somatosensory pRF size for both HV and FHD groups (**Supplementary Figure 3).**

**Figure 6:**
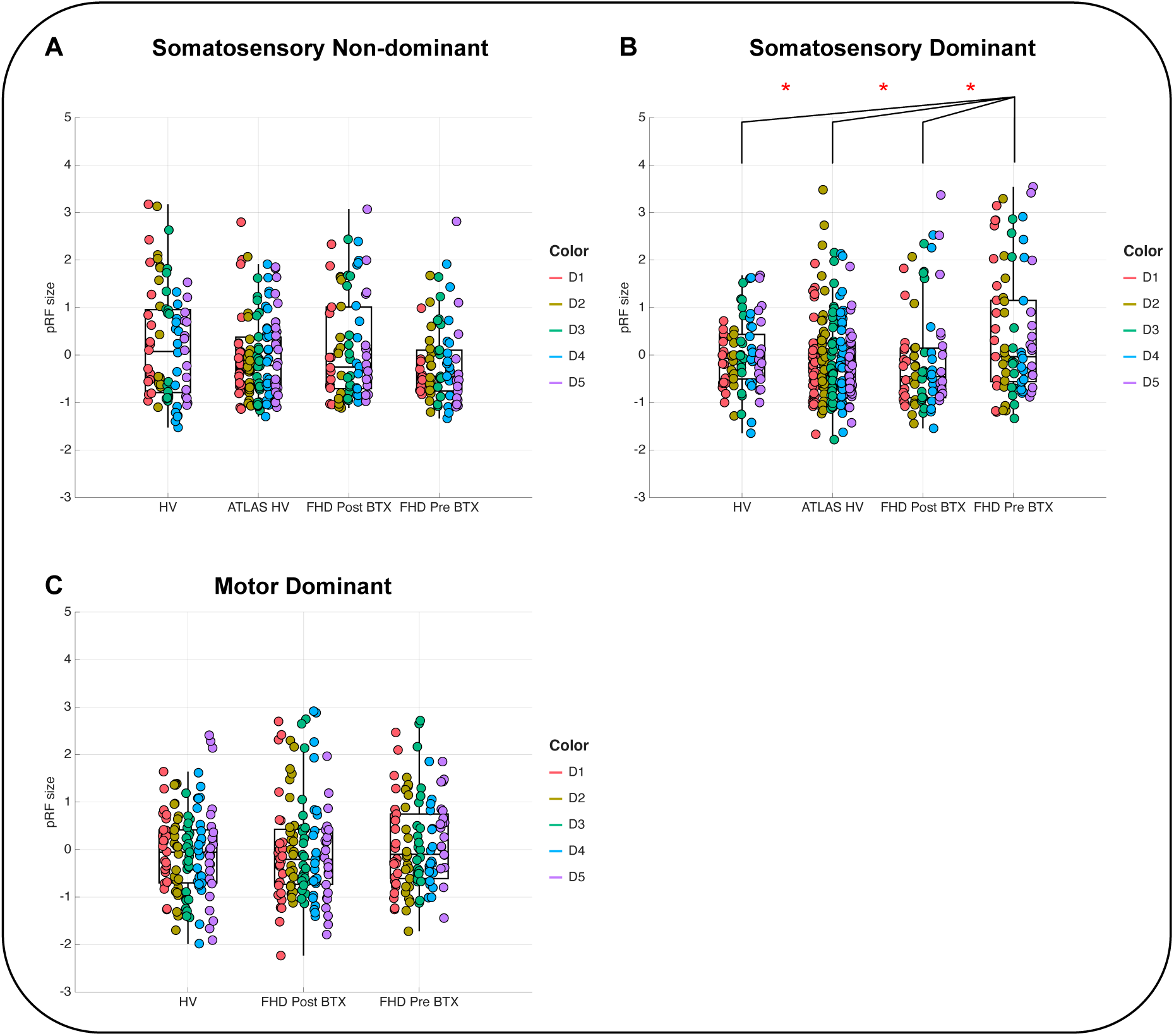
Population receptive field size across all digits, split into the study groups (HV, FHD Post-BTX, and FHD Pre-BTX) as well as the Atlas HVs. A) non-dominant (left) hand and B) dominant (right) hand for the somatosensory task; C) dominant (right) hand for the motor task. * indicates a significant difference (p<0.05), and the height of the bracket indicates which group had the larger pRF size.

**Table 1** and **Figure 7** show the pRF size ordering across digits in each group for non-dominant and dominant hands for the somatosensory task and dominant hand for the motor task. While HV and FHD Post-BTX showed greater pRF sizes for D2,3,4,5 than D1 this was not seen for FHD Pre-BTX in the dominant hand. In the non-dominant hand, all groups had similar pRF size trends, e.g. D2,3,4 > D1. **Table 1** also shows pRF results separated by Brodmann areas, with BA2 larger than BA3a and 3b in all groups in the dominant hand, however, in the non-dominant hand, only Post-BTX and Atlas groups show BA2 > BA3b.

**Figure 7:**
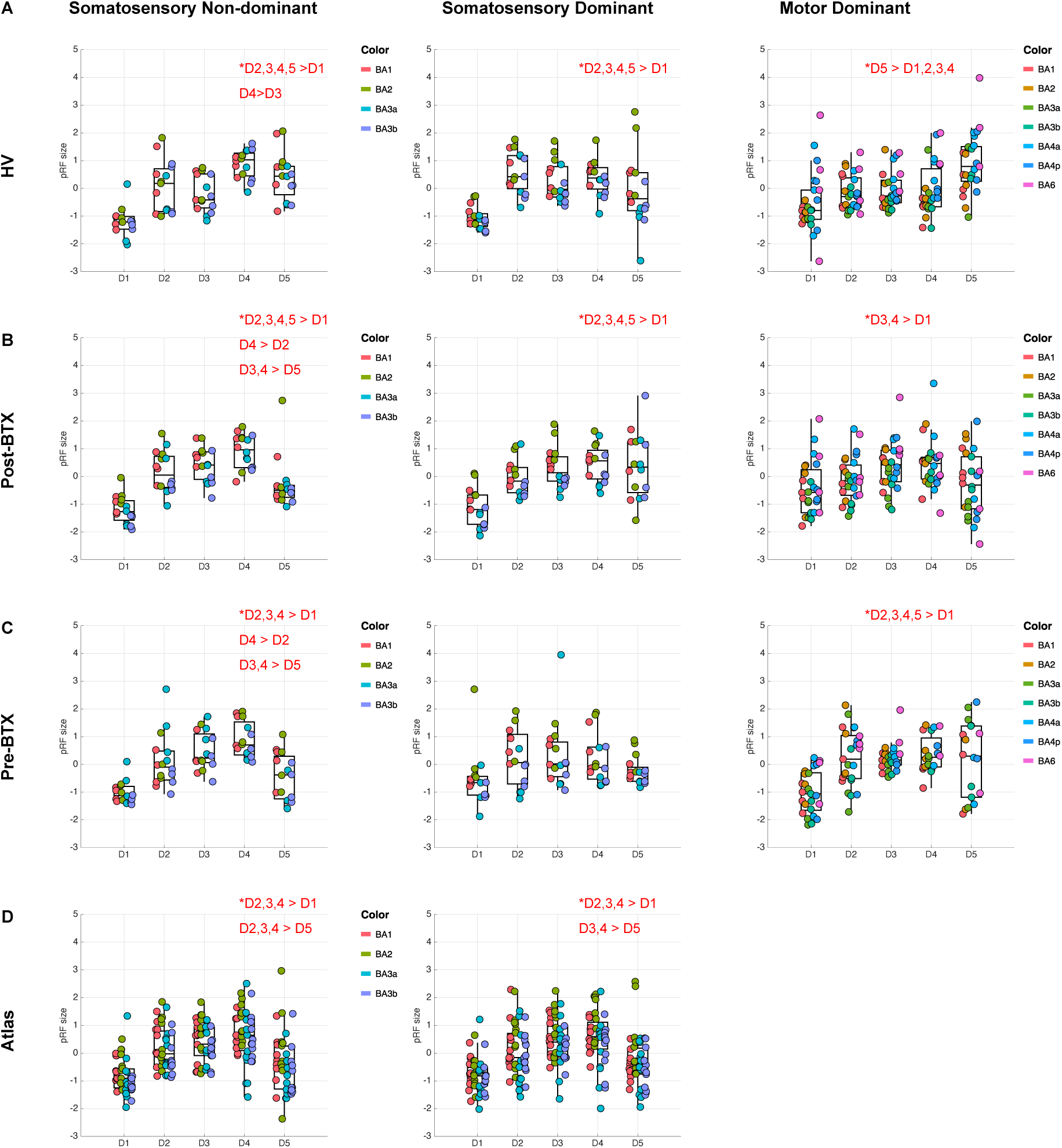
Box plots of population receptive field size, with z-scores shown for each digit (x-axis) and coloured by Brodmann area. Each row is a different group (HV, FHD Post-BTX, FHD Pre-BTX, Atlas HVs). Red text on each plot indicates significant differences between digits. Left column is non-dominant (left) hand and middle column is dominant (right) hand for the somatosensory task; and right column is dominant (right) hand for the motor task.

**Table 1:** Summary of significant differences in pRF size by digit and Brodmann area.

|  | pRF by digits |  |  | pRF by Brodmann area |  |  |
| --- | --- | --- | --- | --- | --- | --- |
|  | <u>Somato Non-Dom</u> | <u>Somato Dom</u> | <u>Motor</u> | <u>Somato Non-Dom</u> | <u>Somato Dom</u> | <u>Motor</u> |
| **HV** | D2,3,4,5>D1<br>D4>D3 | D2,3,4,5>D1 | D5>D1,2,3,4 | No difference | BA2>BA3a&3b | BA6>BA3a,3b&1 |
| **FHD Post-BTX** | D2,3,4,5>D1<br>D4>D2<br>D3,4 >D5 | D2,3,4,5>D1 | D3,4>D1 | BA2>BA3b | BA2>BA3a | BA2>BA3a<br>BA4a>BA3a |
| **FHD Pre-BTX** | D2,3,4>D1.<br>D4>D2.<br>D3, 4 > D5 | No difference | D2,3,4,5>D1 | No difference | BA2>BA3a&3b | No difference |
| **Atlas** | D2,3,4>D1<br>D2,3,4>D5 | D2,3,4>D1<br>D3, 4>D5 | Not acquired | BA2>BA3a&3b | BA2>BA3a&3b | Not acquired |

### 3.5 Resting state fMRI

There was a significant (p<0.001, uncorrected) difference in seed-based connectivity between the seed in left postcentral gyrus (contralateral to dominant hand) and projection to right postcentral gyrus, with lower connectivity in FHD, **Figure 8**. There was no difference in FC in FHD between Post-BTX and Pre-BTX visits.

**Figure 8:**
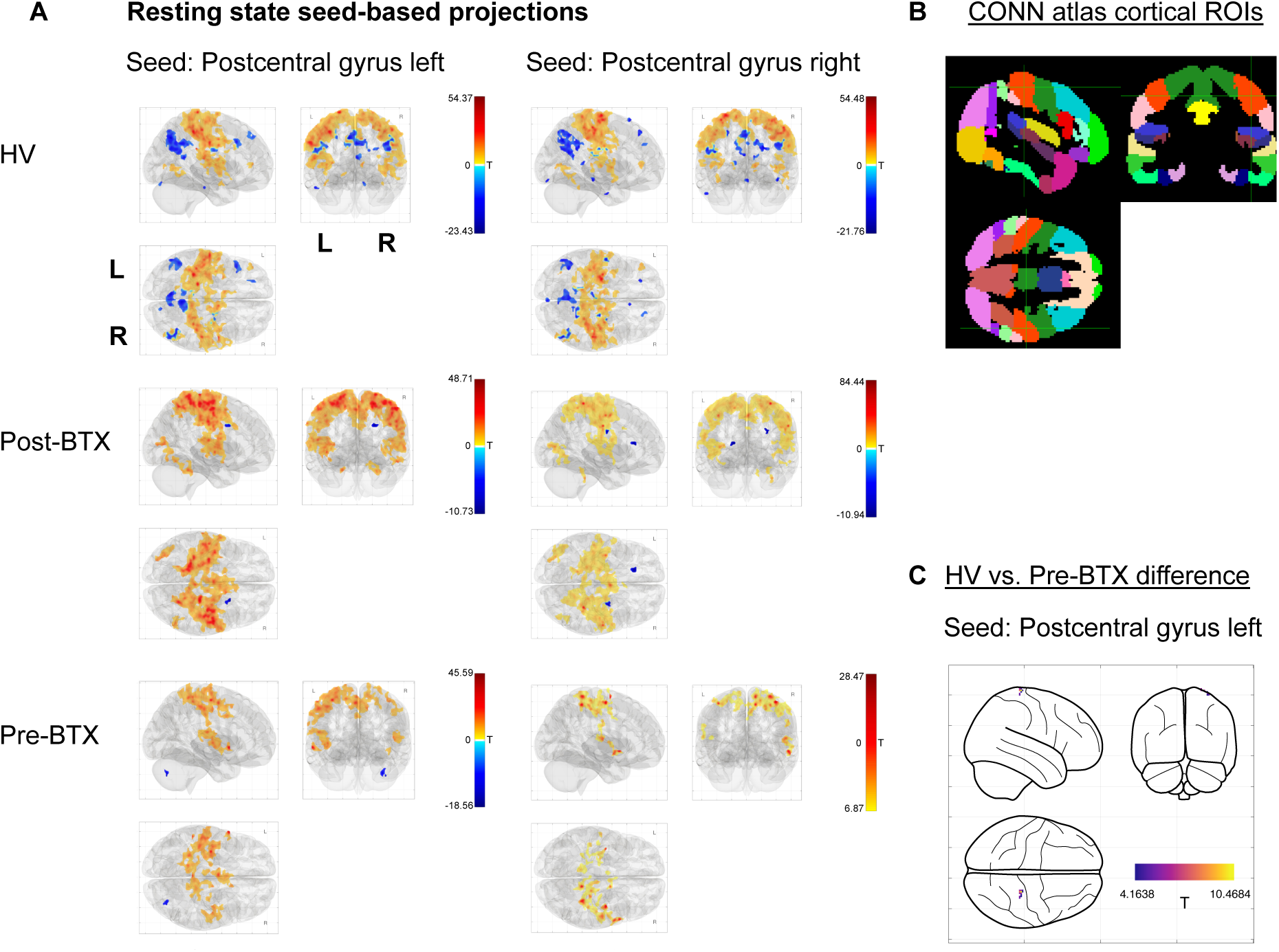
A) Resting state seed-based functional connectivity analysis. Seeds placed in post-central gyrus, and the projection maps for each group are shown. B) CONN cortical ROIs based on Harvard Oxford parcellation are shown. The green crosshair marks the postcentral gyrus. C) Significant difference between groups for a seed in the left (contralateral to dominant hand) postcentral gyrus projecting to right postcentral gyrus. This indicates reduced FC in FHD.

## 4 DISCUSSION

This work has assessed behavioural and functional measures of digit localisation and population receptive field (pRF) size in Focal Hand Dystonia (FHD) patients compared to healthy volunteers, and the effect of treatment with Botulinum Toxin (BTX) using high spatial resolution fMRI.

### 4.1 Behavioural temporal and spatial thresholds in FHD

The spatial acuity threshold measured using the grating orientation task (GOT) was larger in FHD patients compared to healthy controls, supporting the hypothesis of reduced spatial acuity in FHD (28). In contrast, temporal-based temporal discrimination task (TDT) measures did not show any significant differences between groups, this may be due to large variability in the ‘TDT’ measure which was consistent with previous literature (35).(Mikkelsen et al., 2020)

### 4.2 Dominant (affected) and Non-Dominant (unaffected) hand digit maps in FHD

Digit maps on the flattened cortical surface showed clear digit-specific activity patterns for both the travelling wave (TW) phase-analysis and general linear model (GLM) winner-takes-all maps in healthy volunteers (HV) and FHD patients. These findings contest early work showing widespread digit disruption in FHD (2,45) but supports more recent work that have shown intact digit representation within primary sensorimotor cortex of individuals with musician’s dystonia (15). There were however significantly lower Dice coefficients in the dominant affected hand of the FHD group compared to the HV group, with the HV group better matching the probabilistic atlas; this points to a degree of disorganisation or increased variability in the digit representations in the affected hand likely due to abnormal sensorimotor integration (28). It should be noted that early functional magnetic resonance imaging (fMRI) studies (2) were performed at coarser spatial resolution and assessed digit maps in un-flattened space based on Euclidean distances between digits, so differences may have arisen due to folding structure or limitations of spatial resolution or altered amplitude of responses in dystonia patients previously.

It was also shown that the dominant hand had larger digit regions of interest (ROIs) than non-dominant hand for both FHD and HV groups. Digit 5 (D5) was significantly smaller than all other ROIs, agreeing with literature on this digit having the least representation in terms of size, for an overview of cortical magnification of digits, see (46).

For all groups there was large overlap in the digit maps in postcentral gyrus for the somatosensory and motor tasks as evidenced by Dice coefficients. This overlap reflects the anatomical and functional relationship of primary S1 and M1, which form an integrated sensorimotor representation of the digits (47).

### 4.3 Comparison of central tendency scores of digit maps to the probabilistic atlas

The central tendency matrices showed good comparison to the atlas for HV and FHD patients. However, in FHD patients the dominant hand had a significantly lower central tendency than the non-dominant hand. This suggests that in FHD, the dominant hand has larger less centrally matched digit representations whilst the non-dominant hand has smaller more centrally represented digits.

It is noted that the differences between groups may be due to atypical HVs, however, when comparing groups to the Atlas, the HVs perform equal to or better than FHD patients (**Figure 5B).**

### 4.3 Population receptive field sizes in FHD

For the dominant hand, pRF sizes were significantly larger for the FHD Pre-BTX group compared to the HV group and the FHD Post-BTX group. Smaller pRF sizes indicate better tactile acuity, while larger pRFs indicate reduced sensitivity to spatial frequency. This suggests that BTX treatment reduces pRF size to a similar level to HVs, but this increases again ∼3 months later. This finding is consistent with lower Dice scores in the dominant hand, identifying subtle disruption in the dominant affected hand at the BTX visit.

pRF analysis showed that in all groups and hands, the thumb had the smallest pRF size (D2, D3, D4, D5 > D1). This ordering is consistent with electrophysiological and primate literature, whereby D1 has the highest receptor density, smallest receptive fields compared to the other digit tips and the largest cortical magnification (48,49).

Previous fMRI pRF studies have also shown D5 to be larger than the other digits. Interestingly, we showed in some cases that the pRF size of D3 and D4 was greater than D5, where other studies have found D5 to have the largest pRF density size. This may be due to fewer good fits of the pRF fitting algorithm at D5 (46) as less volume is dedicated to D5 in the somatosensory hand map (50).

When comparing pRF size between the dominant and non-dominant hands there was no significant difference (**Supplementary Figure 3)**.

Our finding of no significant difference between dominant and non-dominant hands in digit representations and pRF size in our healthy participants agrees with previous work which showed no significant differences in volume, activation strength, and Euclidean distance between hands (51), or tactile acuity (52).

Motor pRF sizes, measured across both post-central and pre-central gyrus in response to the motor task, were larger than somatosensory pRF size. This is consistent with previous literature which showed larger pRF sizes in solely precentral gyrus (53), and electrophysiologically where complex motor movements have been reported (54).

### 4.5 Resting state functional connectivity in FHD

Resting state data identified lower functional connectivity (FC) in FHD as compared to the HV group, but no differences between FHD Post-BTX and Pre-BTX groups. This is consistent with previous literature, which shows lower FC in sensorimotor regions as compared to controls at rest (22,55) in more participants (n=16, 15, respectively), and the lower sample size here can explain the fewer differences between groups seen here.

### 4.6 Limitations

This study had a relatively small sample size for FHD patients but suggests alterations in the dominant (affected) hand. Future studies should build on this work with a larger study in FHD patients. In future it would also be of interest to compare motor task responses in both the Dominant and non-Dominant hand.

In conclusion, spatial acuity was raised in FHD patients compared to healthy controls. The FHD group had some alterations in their Dominant affected hand with both lower Dice scores and larger pRF sizes between Post-BTX and Pre-BTX. These results provide some evidence to support a neuronal change in the cortical representation of the affected hand in focal hand dystonia and that BTX treatment in FHD patients can alter pRF size in the dominant affected hand.

## Data availability statement

The raw data supporting the conclusions of this article will be made available by the authors on request.

## Ethics statement

The study was conducted with the approval of the University of Nottingham Faculty of Medicine and Health Sciences Ethics Committee for healthy volunteers and of the Health Research Authority Research Ethics Committee (REC 17/EM/0368) for FHD patients. All participants gave written informed consent. The studies were conducted in accordance with the local legislation and institutional requirements. The participants provided their written informed consent to participate in this study.

## Author contributions

MA: study design, data collection, data analysis, manuscript draft. RS: study design, data collection, data analysis. PG: data collection, manuscript editor. DP: data collection, manuscript editor. GON: data collection, manuscript editor. DS: data analysis, manuscript editor. MH: data collection, manuscript editor. SF: study design, data collection, data analysis, manuscript draft. All authors contributed to the article and approved the submitted version.

## Non-standard abbreviations

AMP: Amplitude Thresholding Task
BOLD: Blood Oxygenation Level Dependent contrast
BTX: Botulinum toxin
CT: Central Tendency
FC: Functional Connectivity
FHD: Focal Hand Dystonia
fMRI: Functional magnetic resonance imaging
FoM: Figure of Merit
GOT: Grating Orientation Task
Post-BTX: Focal Hand Dystonia patients ∼30 days after Botulinum toxin treatment
Pre-BTX: Focal Hand Dystonia patients ∼100 days after Botulinum toxin treatment
pRF: Population Receptive Field
rs-fMRI: Resting State fMRI
TDT: Temporal Discrimination Task

## Acknowledgements

We thank the focal hand dystonia participants and healthy volunteers who took part in the study.

## Generative AI statement

The author(s) declare that no Generative AI was used in the creation of this manuscript.

## Contribution to the Field Statement – not for in draft to upload on website on submission

Focal hand dystonia is a neurological condition that affects the digits and can be treated with botulinum toxin (BoNT-A) injections to relieve muscle spasm. This treatment is symptomatic, and it is unclear how it affects brain areas responsible for digit movement and sensation.

We used high-resolution functional magnetic resonance imaging to assess whether digit representations differ between healthy people and individuals with focal hand dystonia. Patients were scanned at two timepoints: (1) shortly after BoNT-A treatment, when its effects were strongest, and (2) three months later, when the effects had worn off.

Patients’ digit maps showed the typical digit ordering overall and compared well to a normative probabilistic atlas. However, we detected subtle differences in FHD patients: (1) The dominant affected hand maps in individuals with FHD showed more spatial disorder in digit to healthy controls. (1) Sensory receptive sizes in the dominant affected hand were smaller shortly after BoNT-A treatment, then enlarged again after three months, suggesting the treatment may temporarily alter brain organization.

These findings add to our understanding of how digits are represented in hand dystonia, and how they respond to treatment.

---

**Supplementary Figure 1.**
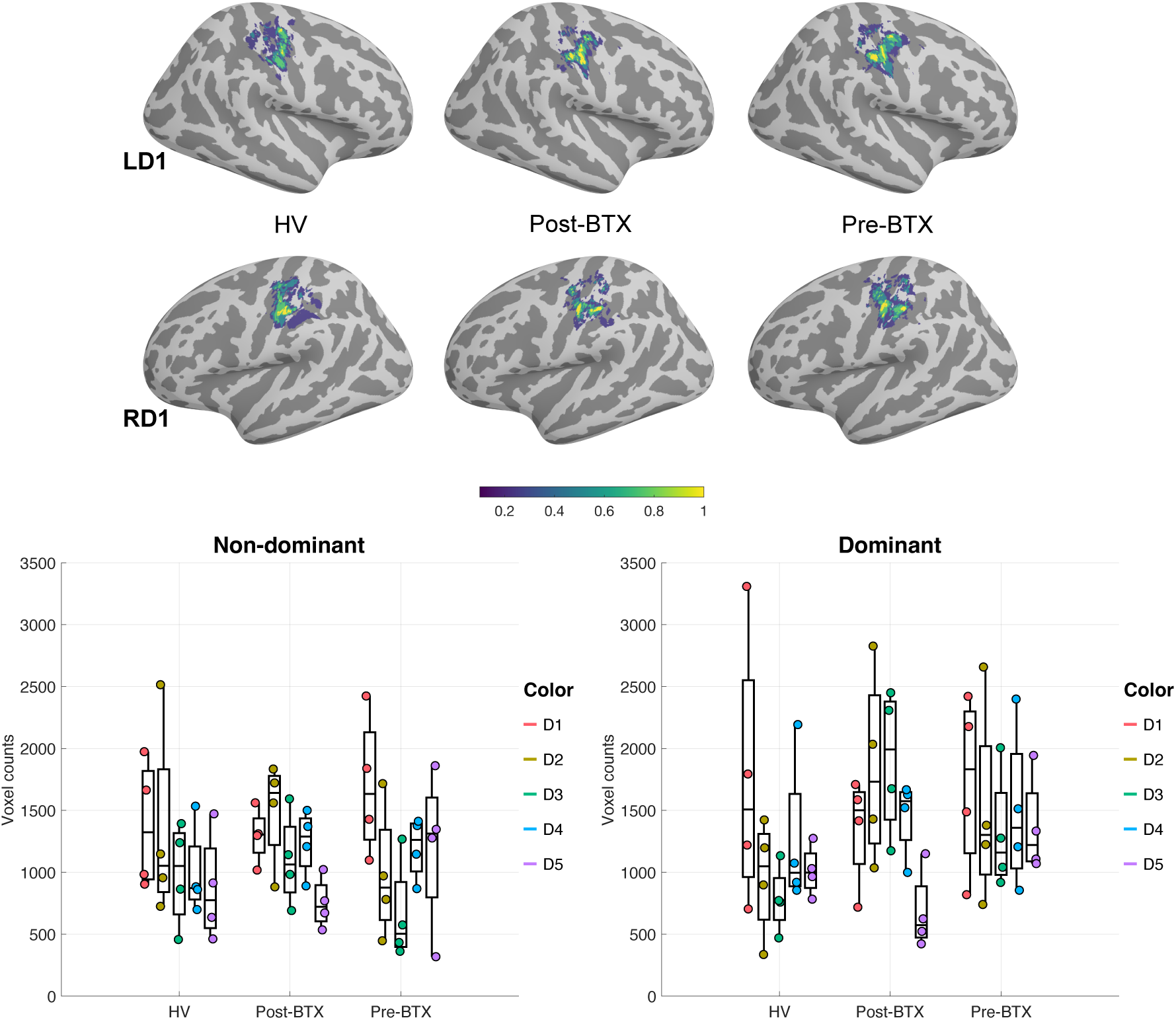
ROIs are shown (RD1 and LD1) for each group. Voxel counts are shown for each group (HV, BTX, NoBTX), for left) non-dominant and right) dominant hands. Each data point corresponds to a single participant’s digit ROI in fsaverage space.The dominant hand was larger than the non-dominant hand across groups. D5 was significantly smaller in all groups.

**Supplementary Figure 2.**
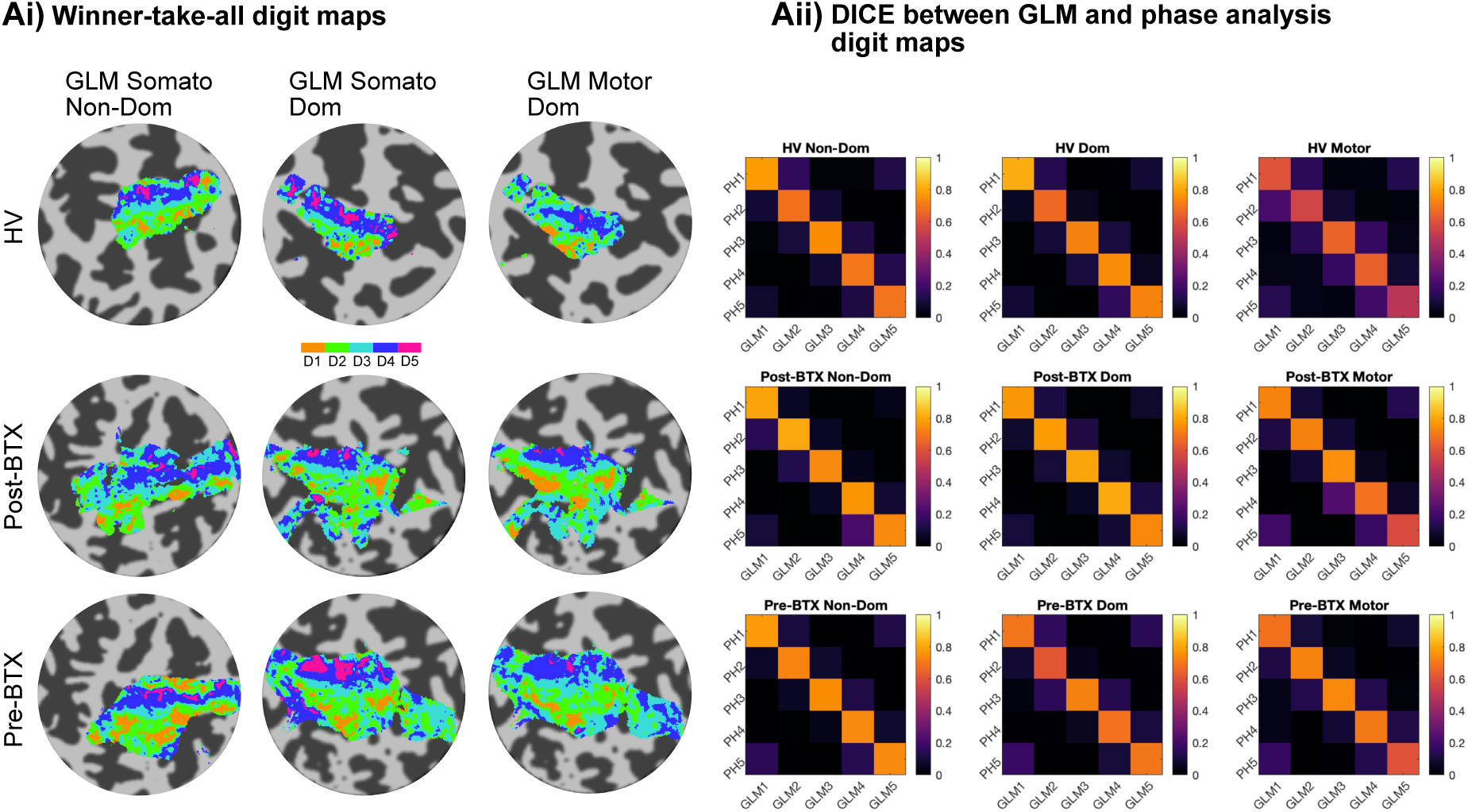
Figure A) GLM analysis of travelling wave fMRI data for digit localisation. i) “winner-take-all” maps shown for an example participant from each group. ii) DICE coefficient matrices showing excellent correspondence between the travelling wave GLM and phase analysis (‘PH’) for digit localisation.

**Supplementary Figure 3.**
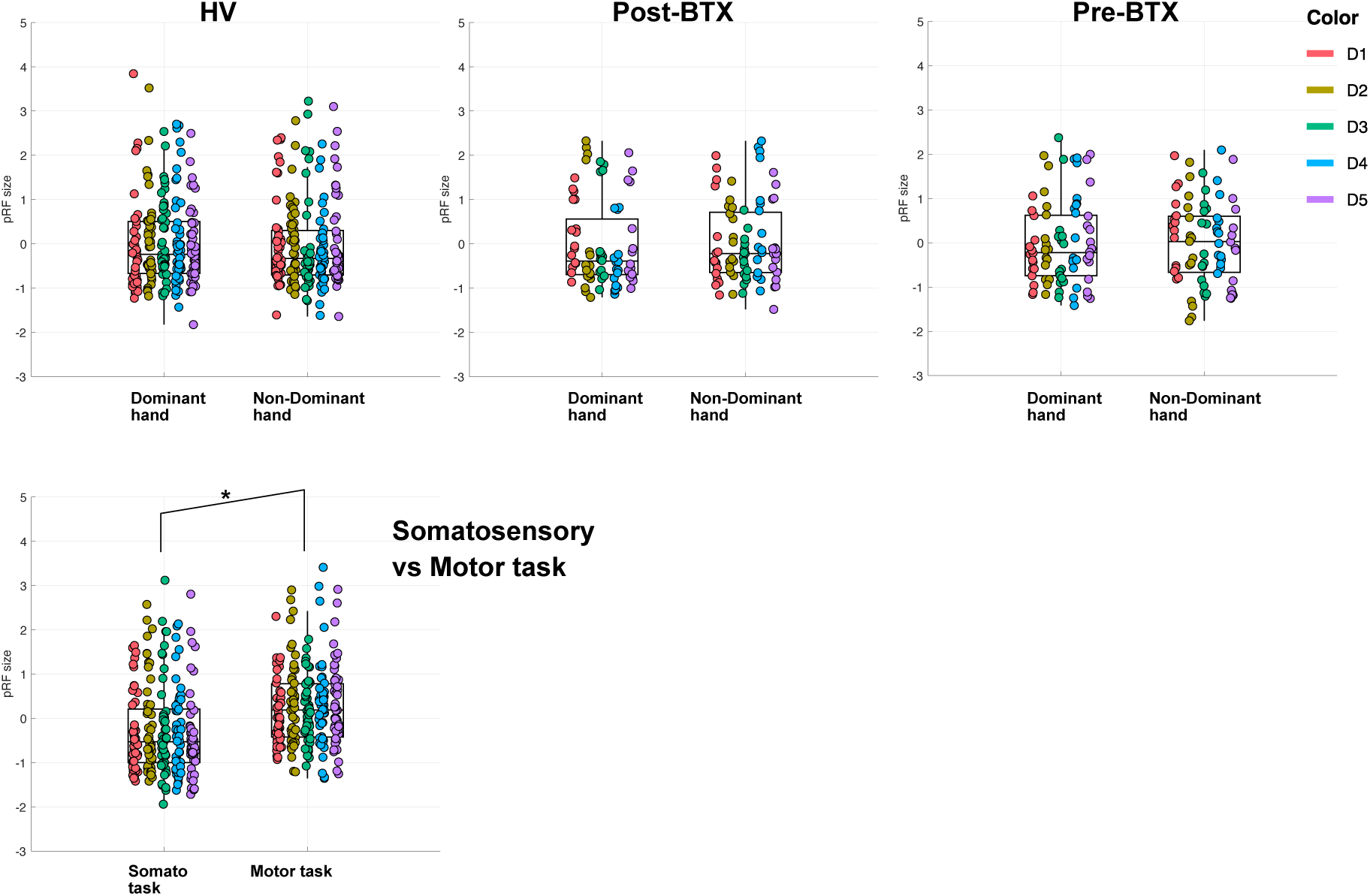
pRF size separated by hand dominance, for each group. Motor pRFs across groups were significantly larger than somatosensory pRFs.

**Supplementary Figure 4.**
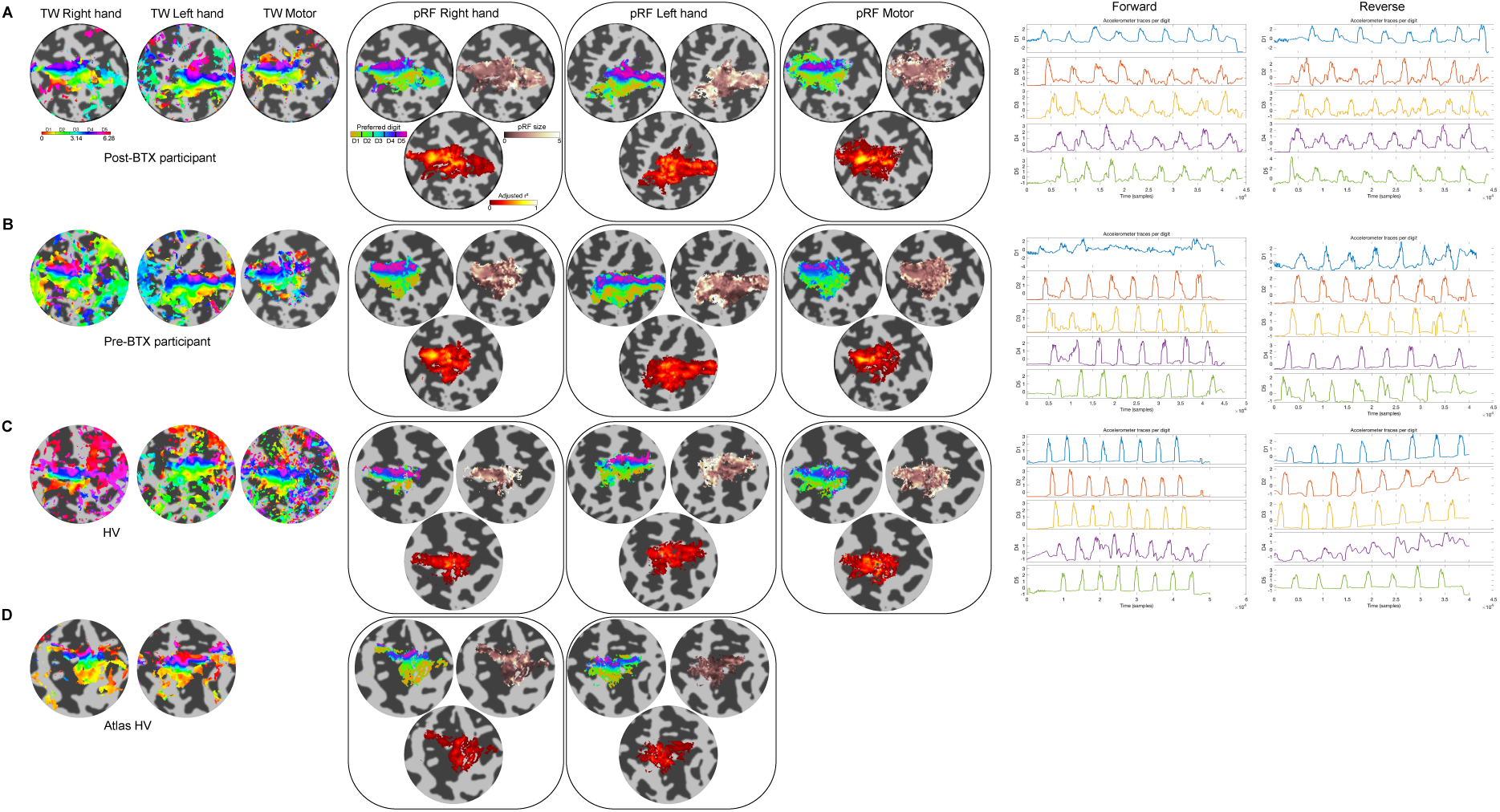
Travelling wave and population receptive field flat maps for a single example participant from each group 1) post-Botox; 2) pre-Botox; 3) healthy volunteers; 4) atlas healthy volunteer.

