## Supplementary Methods for "Assessing the digit organisation of focal hand dystonia using 7T functional MRI"

**Supplementary Table 1** Within- and between-subject coefficient of variation (CV) of behavioural measures defined as a percentage (%) for n=10 healthy volunteer participants.

| Task | Hand | Session | Mean | Std | CV_between | CV_within | CV_between_error | CV_within_error |
|---|---|---|---|---|---|---|---|---|
| Temporal Discrimination | Right | Session 1 | 51 | 36 | 71.4 | 40.0 | 8.2 | 12.2 |
| | | Session 2 | 47 | 35 | 74.4 | | 8.2 | |
| | | All sessions | 49 | 35 | 71.0 | | 5.5 | |
| Temporal Discrimination | Left | Session 1 | 35 | 34 | 94.9 | 46.1 | 10.2 | 11.4 |
| | | Session 2 | 53 | 27 | 51.1 | | 4.4 | |
| | | All sessions | 44 | 31 | 70.2 | | 4.9 | |
| Temporal Discrimination Brain Gauge | Right | Session 1 | 35 | 18 | 51.4 | 29.9 | 3.0 | 6.3 |
| | | Session 2 | 31 | 11 | 37.3 | | 1.3 | |
| | | All sessions | 33 | 15 | 45.4 | | 1.5 | |
| Temporal Discrimination Brain Gauge | Left | Session 1 | 26 | 24 | 92.6 | 31.9 | 7.1 | 8.0 |
| | | Session 2 | 21 | 6 | 29.1 | | 0.6 | |
| | | All sessions | 24 | 17 | 74.1 | | 2.9 | |
| Amplitude discrimination 30Hz | Right | Session 1 | 4.2 | 2.1 | 49.8 | 21.1 | 0.3 | 6.2 |
| | | Session 2 | 4.3 | 2.0 | 47.3 | | 0.3 | |
| | | All sessions | 4.2 | 2.0 | 47.2 | | 0.2 | |
| Amplitude discrimination 30Hz | Left | Session 1 | 4.4 | 2.9 | 66.8 | 28.0 | 0.6 | 7.7 |
| | | Session 2 | 5.6 | 4.0 | 72.0 | | 0.9 | |
| | | All sessions | 5.0 | 3.5 | 69.9 | | 0.5 | |
| Amplitude discrimination 200Hz | Right | Session 1 | 3.7 | 1.6 | 44.7 | 34.4 | 0.2 | 5.1 |
| | | Session 2 | 5.0 | 3.0 | 59.1 | | 0.6 | |
| | | All sessions | 4.3 | 2.4 | 56.3 | | 0.3 | |
| Amplitude discrimination 200Hz | Left | Session 1 | 4.3 | 2.4 | 56.6 | 43.3 | 0.4 | 11.8 |
| | | Session 2 | 4.7 | 3.3 | 70.8 | | 0.8 | |
| | | All sessions | 4.5 | 2.9 | 63.2 | | 0.4 | |
| Grating orientation task | Right | Session 1 | 1.7 | 0.5 | 30.3 | 6.2 | 0.05 | 1.7 |
| | | Session 2 | 1.6 | 0.6 | 36.8 | | 0.07 | |
| | | All sessions | 1.7 | 0.6 | 32.8 | | 0.04 | |

**Supplementary Methods: Assessing digit correspondence of TW and General Linear Model (GLM) analysis**

Somatotopy of both hands and motortopy data of the dominant hand were analysed using a GLM to assess in each group how GLM digit maps compare to phase-analysis digit maps.

For the somatosensory task, a 4s "ON" boxcar convolved with the canonical haemodynamic response function (HRF) was used in the GLM to define each stimulation period. For the motortopy task, each digit was modelled in the GLM using the exact timings of movement onset and offset from the accelerometer glove.

Digit location responses to each stimulation period were defined ([1 0 0 0 0]; [0 1 0 0 0]; [0 0 1 0 0]; [0 0 0 1 0]; [0 0 0 0 1]) and using a "winner-take-all" approach of the highest beta weight, each voxel was assigned to a given digit. The GLM "winner-take-all" approach was compared to the Fourier analysis of digit localisation.
